# The Nuclear Lamina Binds the EBV Genome During Latency and Regulates Viral Gene Expression

**DOI:** 10.1101/2021.08.05.455214

**Authors:** Lisa Beatrice Caruso, Rui Guo, Kelsey Keith, Jozef Madzo, Davide Maestri, Sarah Boyle, Jason Wasserman, Andrew Kossenkov, Benjamin E. Gewurz, Italo Tempera

## Abstract

The Epstein Barr virus (EBV) infects almost 95% of the population worldwide. While typically asymptomatic, EBV latent infection is associated with several malignancies of epithelial and lymphoid origin in immunocompromised individuals. In latently infected cells, the EBV genome persists as a chromatinized episome that expresses a limited set of viral genes in different patterns, referred to as latency types, which coincide with varying stages of infection and various malignancies. We have previously demonstrated that latency types correlate with differences in the composition and structure of the EBV episome. Several cellular factors, including the nuclear lamina, regulate chromatin composition and architecture. While the interaction of the viral genome with the nuclear lamina has been studied in the context of EBV lytic reactivation, the role of the nuclear lamina in controlling EBV latency has not been investigated. Here, we report that the nuclear lamina is an essential epigenetic regulator of the EBV episome. We observed that in B cells, EBV infection affects the composition of the nuclear lamina by inducing the expression of lamin A/C, but only in EBV+ cells expressing the Type III latency program. Using ChIP-Seq, we determined that lamin B1 and lamin A/C bind the EBV genome, and their binding correlates with deposition of the histone repressive mark H3K9me2. By RNA-Seq, we observed that knock-out of lamin A/C in B cells alters EBV gene expression. Our data indicate that the interaction between lamins and the EBV episome contributes to the epigenetic control of viral gene expression during latency, suggesting a restrictive function of the nuclear lamina as part of the host response against viral DNA entry into the nucleus.

**AUTHOR SUMMARY:** Epstein-Barr virus (EBV) is a common herpesvirus that establishes a lifelong latent infection in a small fraction of B cells of the infected individuals. In most cases, EBV infection is asymptomatic; however, especially in the context of immune suppression, EBV latent infection is associated with several malignancies. In EBV+ cancer cells, latent viral gene expression plays an essential role in sustaining the cancer phenotype. We and others have established that epigenetic modifications of the viral genome are critical to regulating EBV gene expression during latency. Understanding how the EBV genome is epigenetically regulated during latent infection may help identify new specific therapeutic targets for treating EBV+ malignancies. The nuclear lamina is involved in regulating the composition and structure of the cellular chromatin. In the present study, we determined that the nuclear lamina binds the EBV genome during latency, influencing viral gene expression. Depleting one component of the nuclear lamina, lamin A/C, increased the expression of latent EBV genes associated with cellular proliferation, indicating that the binding of the nuclear lamina with the viral genome is essential to control viral gene expression in infected cells. Our data show for the first time that the nuclear lamina may be involved in the cellular response against EBV infection by restricting viral gene expression.

## INTRODUCTION

Epstein Barr Virus (EBV) is a human gamma-herpesvirus that establishes lifelong latent infection in almost 95% of the population [1–4]. EBV infection is generally asymptomatic; however, EBV latency is associated with several malignancies, including Burkitt lymphoma, Hodgkin’s lymphoma, gastric cancer, and nasopharyngeal carcinoma [5–9] . During latent infection, EBV persists as multicopy nuclear episomes [10, 11], and viral gene expression is restricted to a few latent genes. These latent genes are expressed in different patterns — referred to as latency types — that induce distinct responses in the infected cells and are associated with different types of malignancies [12, 13]. Type I latency is characterized by the expression of the viral protein EBNA1 [14], which is necessary for viral DNA replication and episome maintenance by tethering the EBV genome to the host chromosomes [10, 15]. Type III latency is associated with the expression of all the EBV latent proteins: six nuclear antigens (EBNA1, EBNA2, EBNA3A, EBNA3B, EBNA3C, and EBNA-LP) and three membrane proteins (LMP1, LMP2A, and LMP2B), in addition to a few non-coding RNAs [12]. Switching between these different latency types is an essential strategy for EBV to persist in the host and evade immune detection; however, the mechanisms controlling EBV latency are only partially understood.

We and others have previously reported differences in histone modifications and DNA methylation levels of the EBV genome across different latency types [16–21]. Furthermore, EBV genomes representing distinct latency types have vastly different three-dimensional chromatin conformations that correlate with viral gene expression [22, 23]. Overall, these observations indicate that dynamic changes in chromatin composition and architecture can be critical for initiating and maintaining latency programs, but how these epigenetic programs are established remains unclear.

In eukaryotes, the epigenetic status and conformation of the genome can be influenced by the interaction of genomic regions with the nuclear lamina [24, 25]. The nuclear lamina is a filamentous protein network located at the nuclear periphery that is formed by proteins known as lamins that are grouped into two different families (type A and B lamins) based on sequence homology and molecular weight. B-type lamins (B1 and B2) localize only at the nuclear periphery and are ubiquitously expressed in mammalian cells from two different genes (*LMNB1* and *LMNB2*), while A-type lamins (A and C) are also present in the nuclear interior and are encoded by differential splicing of a single gene (*LMNA*) that is expressed only in differentiated cells [24, 26, 27].

The nuclear lamina interacts with genomic DNA at regions of the chromatin called lamin associated domains, or LADS, ranging in size between 0.1 and 10 Mb, whose spatial location within the nucleus shifts dynamically during cell differentiation [28, 29]. Both lamin A and lamin B can interact with chromatin to form LADS, which are referred to as A-LADS and B-LADS, respectively [30]. While most of the A- and B-LADs overlap, the two lamin types can also form distinct LADs, suggesting that A- and B-type lamins may have complementary and specific functions in organizing the genome architecture within the nuclear space [31–33]. For example, lamin A-LADs tend to be positioned more toward the center of the nucleus than the other types of LADs [30]. When localized at the nuclear periphery, all LADs have low transcription levels and are associated with repressive histone marks, including H3K9me2/3 and H3K27me3 [34–37]. The H3K9me2 histone mark plays an essential role in positioning the LADs at the nuclear periphery, as depletion of the histone methyltransferase G9a, which specifically mono- and demethylates Lysine-9 of histone H3, causes LADs to dissociate from the nuclear lamina [29]. Detachment of the LADs from the nuclear periphery also causes reorganization of the 3D structure of the genome, indicating that the nuclear lamina assists in the spatial organization of the genome [38–40]. These observations suggest that the interplay between LADs and lamins is essential in the dynamic organization of the genome and the regulation of gene expression.

Nuclear lamins play an antiviral role during the replication of herpesviruses, including EBV [41, 42], HSV [43–50], and HCMV [51–56], regulating viral replication and functioning as a physical barrier that prevents the egress of the viral capsid. Moreover, the nuclear lamina interacts with the HSV1 genome and plays a critical role in forming replication compartments during the lytic replication of HSV1 [48]. To counteract the antiviral effect of the nuclear lamina, herpesviruses have evolved lytic genes encoding viral proteins that target the nuclear lamina during a lytic infection. For example, the EBV lytic gene *BGLF4* encodes a viral kinase that interacts with the nuclear lamina and causes its degradation via phosphorylation of lamin A [41]. While the antiviral function of lamins and the nuclear lamina during EBV lytic infection has been described, we still ignore whether the nuclear lamina plays a similar restrictive role during latent viral infection. Based on the ability of lamins to bind the genome and to regulate both the structure and conformation of the LADs, here, we investigated the role of nuclear lamins in the control of the EBV latency states and the potential function of the nuclear lamina as an epigenetic regulator of the different EBV gene expression profiles observed during latency.

We found that infection of B cells with EBV changes the nuclear lamina composition by inducing lamin A/C expression, which is maintained during Type III latency and lost in Type I latency. Consistent with these changes in nuclear lamina composition, we found that lamin B1 binds to several regions of the EBV genome in Type I latency, while lamin A/C binds to a few regulatory viral regions in Type III cells. We observed that the regions of the EBV genome repressed through nuclear lamina binding were enriched for the repressive histone mark H3K9me2. Furthermore, by knocking out *LMNA* expression in lymphoblastoid cell lines (LCLs), we found that lamin A/C regulates the expression of both lytic and latent genes and that the changes in gene expression were associated with changes in the chromatin composition of the EBV genome. Finally, we determined that lamin A/C knock-out has no effects on the 3D organization of the EBV genome, indicating that lamin A/C only influences its chromatin composition.

Altogether, our data indicate that EBV infection affects the nuclear lamina composition by promoting expression of lamin A/C, which contributes to the establishment of specific chromatin landscapes and transcriptional profiles of Type III latent EBV genome.

## RESULTS

### EBV+ B cells adopting the Type III latency program express lamin A/C

The nuclear lamina represents a physical barrier for the nuclear egress of viral capsids and undergoes a dramatic reorganization during herpesvirus lytic infection. However, the effect of EBV latent replication on the nuclear lamina has not been widely explored. To assess whether EBV latent replication alters the nuclear lamina composition, we analyzed the expression of its two major components, lamin B1 and lamin A/C, in a panel of EBV+ B cell lines and primary B cells from a healthy donor (**Figure 1**). Using western blot analysis, we observed that primary B cells and all cell lines express lamin B1. In contrast, lamin A/C expression was present only in EBV+ B cells adopting the Type III latency (Mutu-LCL, B95.8-LCL, GM12878, and Mutu III). This was not due to inherent genetic differences in the host cells or EBV strains, as the lamin A/C protein was differentially expressed in Mutu III cells (Type III) and Mutu I cells (Type I) that are fully isogenic with respect to both the host and EBV genomes.

**Figure 1:**
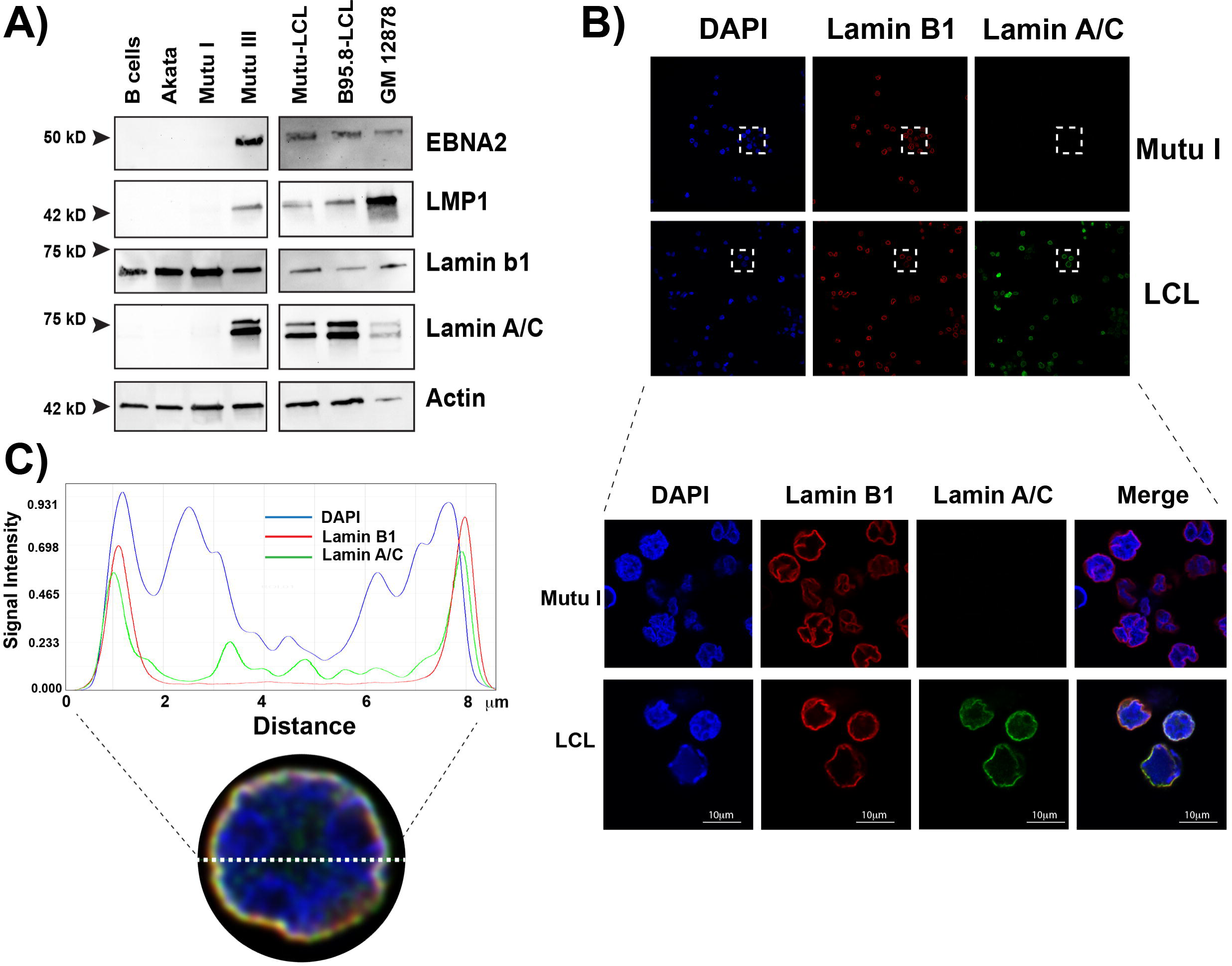
Type I and Type III EBV+ cells express different components of the nuclear lamina. **A)** Western blot for key EBV proteins (EBNA2, LMP1) and nuclear lamina components (lamin B1 and A/C) in EBV-negative cell lines (B cells, Akata) and EBV-positive cell lines that express the type I (Mutu I) or the type III (Mutu III, Mutu-LCL, B95.8-LCK, GM12878) latency program. EBV actin was used as a loading control. Images are representative of at least three independent experiments. **B)** Top: Confocal microscopy analysis of EBV+ Mutu I and LCL cells stained with anti-Lamin B1 and anti-Lamin A/C antibodies and DAPI. The images show that lamin A/C is differentially expressed, depending on the presence of EBV and its latency state. Bottom: Higher magnifications of the boxed areas showing colocalization of the lamin proteins at the nuclear periphery. **C)** Representative quality analysis of the fluorescence intensity in LCL cell nuclei immunostained as described above. The fluorescence intensity was measured along the dotted white line (x-axis) using a Leica software analysis.

To determine whether the differences in the composition of the nuclear lamina correlate with changes in its structure and whether the localization of the lamins differs between EBV latency types, we used immunofluorescence microscopy to assess the specific localization of lamin B1 and lamin A/C in the nucleus of Mutu I cells and LCL cells (Mutu-LCL, Type III) generated by immortalization of primary B cells with the Mutu strain of EBV. We found that lamin B1 was localized at the nuclear rim in both Mutu I and Mutu-LCL cells (**Figure 1B** and **1C**), in agreement with previous observations. In contrast, lamin A/C staining was present only in Mutu-LCL cells, consistent with our western blotting results and confirming that lamin A/C expression is restricted to Type III EBV+ B cells (**Figure 1B**). In Mutu-LCL cells, lamin A/C staining was localized at the nuclear rim (**Figure 1B**) and overlapped with lamin B1 staining (**Figure 1C**). Lamin A/C staining was also present in the nucleoplasm, although with lower intensity than at the nuclear rim, in agreement with previous reports indicating that a fraction of lamin A/C exists in the nucleoplasm (**Figure 1C and 1D**). To further investigate the distribution of lamin A/C and lamin B1 in LCL cells and to validate the presence of lamin A/C in the nucleoplasm, we qualitatively analyzed the fluorescence intensity of lamin B1, lamin A/C and DAPI along the nuclear diameter. We confirmed that the lamin A/C signal was distributed from the nuclear lamina toward the interior of the nucleus, while lamin B1 was localized only at the nuclear lamina (**Figure 1D**). Using laser scanning confocal microscopy in *z*-stack mode, we confirmed that lamin B1 and lamin A/C colocalize at the nuclear rim, but lamin A/C was also present in the interior of the nucleus (**Supplementary video 1 and 2**). Overall, these results show that the composition of the nuclear lamina differs among EBV+ latently infected B cells, with lamin A/C expressed only in cells adopting the Type III latency.

### EBV infection of primary B cells induces lamin A/C expression

Since we observed that primary B cells do not express the lamin A/C protein, we tested whether lamin A/C expression is induced upon EBV infection or as a later consequence of infection (**Figure 2**). First, we analyzed by immunofluorescence microscopy the expression of lamin B1 and lamin A/C in primary B cells from healthy donors before and 24 hours after infection with EBV (B95.8 virus strain). We confirmed that B cells expressed lamin A/C only after EBV infection and that the lamin A/C signal was detected both at the nuclear rim, where it overlapped with lamin B1, and in the nucleoplasm, as observed in Type III EBV+ cell lines (**Figure 2A**). In addition, we observed that the nuclei of infected cells had a lobulated shape (**Figure 2A**). These data indicate that lamin A/C was expressed upon EBV infection, and this change in nuclear lamina composition correlated with altered nuclear shape in infected cells.

**Figure 2:**
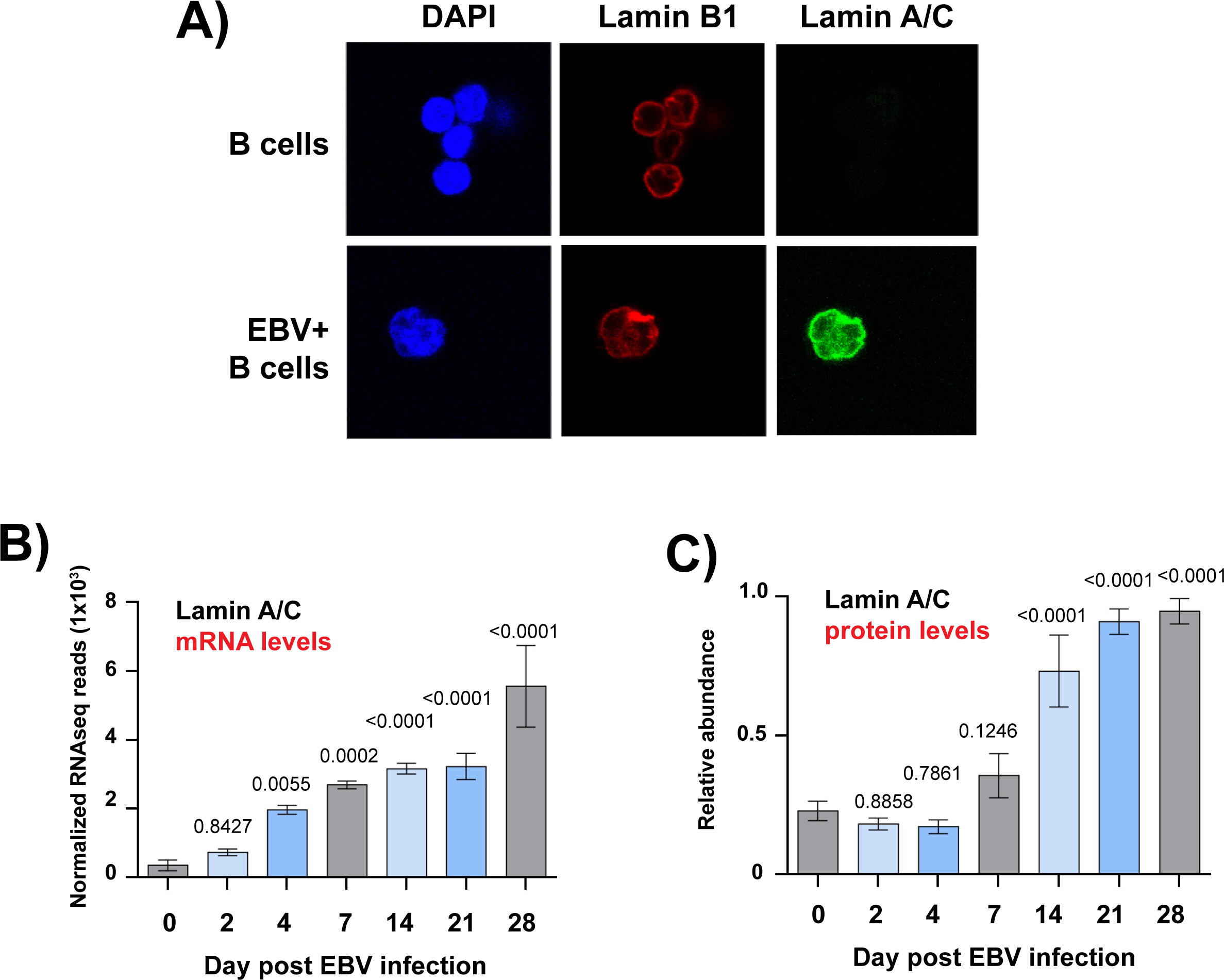
EBV infection of primary B cells induces the expression of lamin A/C. **A)** Immunofluorescence confocal microscopy analysis of B cells from a healthy donor immunostained with lamin B1 (red) and lamin A/C (green) antibodies and DAPI (blu). Top: control (uninfected) cells; bottom: 2 days post-infection with EBV viral particles. **B)** RNA-seq of primary B cells from 3 different healthy donors that were infected with EBV and collected at the indicated time points after viral infection. The plot shows the normalized reads of lamin A/C mRNA. The t test p values for each time point are indicated. **C)** Whole cell proteomic analysis of lamin A/C expression at the indicated time points following infection of primary human B cells as described in B. The plot shows the relative abundance of lamin A/C from 3 biological replicates, representing 12 human donors. The t test p values for each time point are indicated. N=3, Mean ± SD.

It has been reported that both viral and cellular genes are expressed in different patterns during the time necessary for EBV to transform primary B cells to LCLs [57, 58]. We therefore explored whether lamin A/C expression changes during progression of EBV infection by interrogating RNA-Seq datasets of primary B cells from healthy donors before and after EBV infection, at 9 different timepoints (**Figure 2B**). We observed that lamin A/C mRNA abundance significantly increased 4 days post infection and reached a ∼10-fold increase 28 days post infections, when EBV+ B cells exhibit LCL physiology and the Type III latency gene expression program (**Figure 2B**). Using proteomic datasets obtained from human primary B cells before and after EBV infection, we found that lamin A/C protein levels mirrored the surge in lamin A/C mRNA expression, reaching a ∼5-fold increase in the same time frame (**Figure 2C**). Taken together, these results demonstrate that EBV infection and the subsequent establishment of the Type III latency program induce and maintain lamin A/C expression in B cells.

### B cell activation induces lamin A/C expression

In infected B cells, the EBV type III latency program is characterized by the expression of viral latent proteins, including EBNA2 and LMP1, that interfere with B cell biology and mimic key aspects of antigen-mediated B cell activation. We therefore investigated whether physiological activation of B cells also induced lamin A/C expression. To address this question, we compared lamin A/C and lamin B1 expression in primary B cells before and after 24 hours of treatment with interleukin 4 (IL-4), CD40 ligand (CD40L), and CpG oligodeoxynucleotides, a cocktail that mimics antigen activation and stimulates B cell proliferation (**Figure 3**). We achieved ∼65% of activated B cells, as demonstrated by expression of the transmembrane C-Type lectin protein CD69, a marker for B cell activation (Supplementary Figure 1). Analyzing CD69+ B cells and untreated (control) cells by immunofluorescence microscopy, we detected lamin B1 expression at the nuclear rim in both conditions (**Figure 3A**). Lamin A/C expression was observed both at the nuclear rim, colocalizing with lamin B1 signal, and in the nucleoplasm in CD69+ B cells, but not in untreated cells. Furthermore, the nuclei of activated B cells had an irregular and elongated shape (**Figure 3A**). Since the stimulatory cocktail activates both BCR and CD40 signaling pathways, we next dissected which of the B cell stimuli included in the cocktail treatment triggered lamin A/C expression. We used RNA-Seq to quantify the expression of lamin A/C in B cells after treatment with CD40L, CpG, αIgM, or IL-4, individually or in different combinations. B cell activation with CD40L alone was sufficient to induce a 3-fold increase in lamin A/C expression compared to untreated cells, whereas treatment with CpG, αIgM or IL-4 alone had a very modest effect (**Figure 3B**). The increase in lamin A/C expression was similar after treatment with CD40L alone and in combination with the other stimulatory compounds, negating the existence of additive or synergistic effects between CD40 signaling and BCR signaling in inducing lamin A/C expression (**Figure 3B**). Proteomic analysis of lamin A/C expression in B cells before and after the stimulatory treatment described above showed that CD40L was sufficient to induce a significant increase of lamin A/C protein levels compared to control (**Figure 3C**). Consistent with RNA-Seq data, there was no additive or synergistic effect on lamin A/C expression when B cells were treated with CD40L in combination with the other stimulatory compounds (**Figure 3C**). These results indicated that induction of lamin A/C expression is part of the cascade of cellular changes that occur in B cells after activation via T-cell CD40L stimulation.

**Figure 3:**
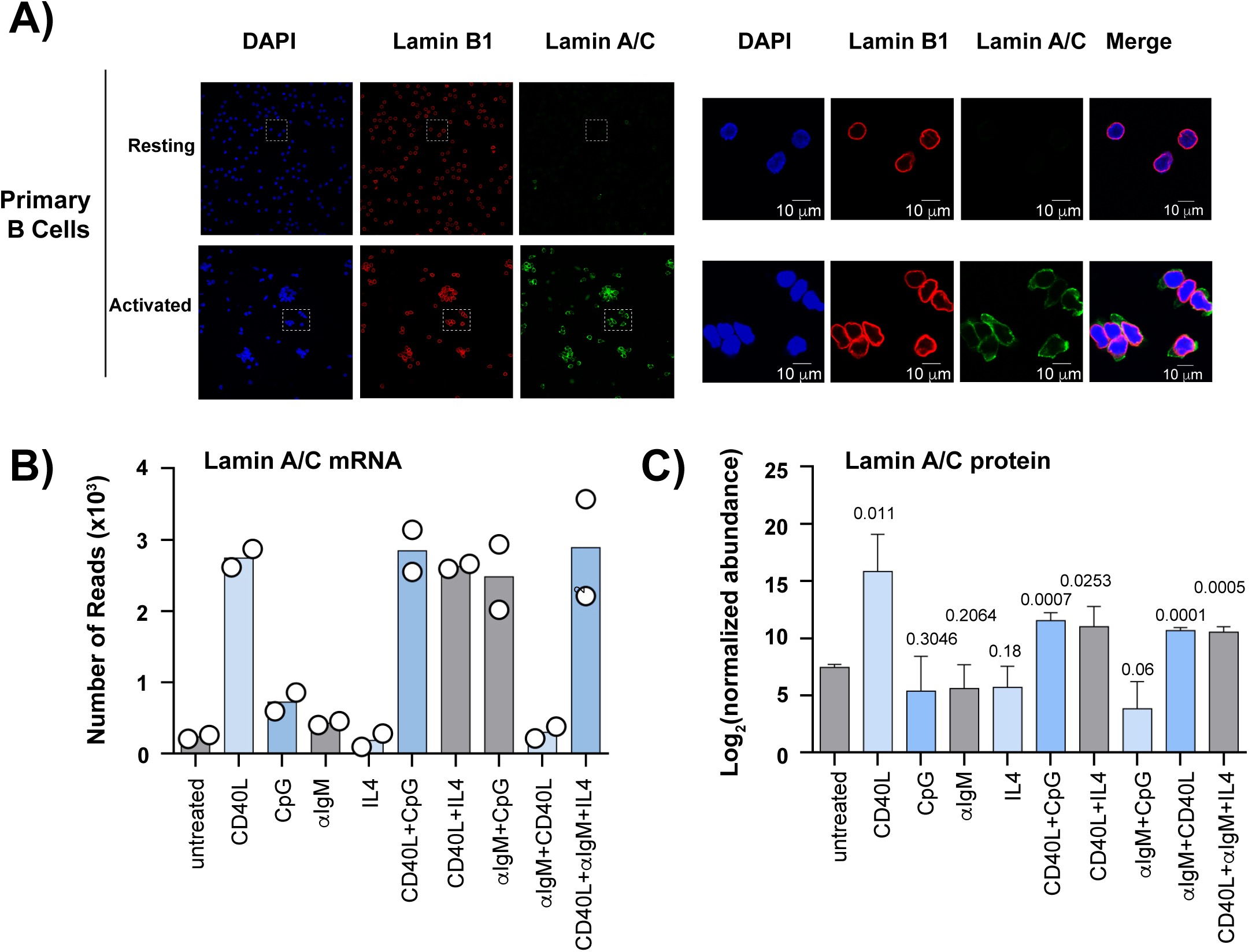
Activation of B cells by CD40 treatment induces lamin A/C expression. **A)** Confocal analysis of resting (top) and activated (bottom) primary B cells from a healthy donor, immunostained with lamin A/C (green) and lamin B1 (red) antibodies and DAPI (blue). For activation, cells were treated for 24 hours with a stimulatory cocktail containing 20 ng/mL interleukin 4 (IL-4), 5uM CD40 ligand (CD40L), and 25 ng/mL CpG oligodeoxynucleotides. The right panel shows higher magnification of the boxed areas in the left panel to show colocalization of the lamin proteins at the nuclear periphery. **B)** RNA-seq of primary B cells from 2 donors. The plot shows the normalized reads for lamin A/C mRNA after a 24-hour stimulation with the indicated ligands (control = untreated cells). The values of two different experiments performed in parallel is shown. **C**) Proteomic analysis of lamin A/C expression in B cells treated as described above with the indicated stimulatory compounds (control = untreated cells). Data are presented as Mean ± SD (N=3). The t test p values for each time point are indicated.

### The EBV genome interacts with the nuclear lamina at viral LADs during latency

The binding of lamin B and lamin A/C to genomic DNA affects the epigenetic modifications of the interacting chromatin regions. Based on the observed changes in nuclear lamina composition between EBV latency types and the increased expression of lamin A/C resulting from EBV infection, we investigated whether the EBV genome interacts with the nuclear lamina during latency and whether this interaction is latency type-specific. We used chromatin immunoprecipitation followed by next-generation sequencing (ChIP-Seq) to assess lamin B1 and lamin A/C binding across the EBV genome in Mutu I (Type I) and LCL (Type III) cells (**Figure 4**). Using a computational peak calling method, we identified areas in the EBV genome enriched for the biding of nuclear lamins in Type I and Type III latency, suggestive of the existence of LADs across the viral latent genome (**Figure 4**, and **Supplementary table 1**). In Type I cells, we identified 23 viral regions enriched for lamin B1, including 3 prominent peaks, 2 of which localized at lytic origins of DNA replication (OriLyt-left and RPMS1-OriLyt), and 1 at the tandemly arranged terminal repeats (TR) region (**Figure 4**, blue track, and **Supplementary table 1**). In Mutu I cells, the width of lamin B1-LADs varied, with an average of 2.4 kbp (**Supplementary table 1**). In contrast, in Type III cells we identified only 3 viral regions associated with lamin B1, coinciding with the three prominent peaks present in Type I cells. (**Figure 4**, green track, and **Supplementary table 1**). In addition to being less numerous, we noticed that lamin B1-LADs were narrower in LCL cells compared to Mutu I. In Type III cells, we identified 16 EBV regions bound by lamin A/C, including a peak at the TR region and a cluster of peaks spanning the region between 110 kbp and 130 kbp (**Figure 4**, orange track, and **Supplementary table 1**). By quantitative ChIP, we confirmed that lamin B1 was bound to the OriLyt-left and TR regions in both Mutu I and LCL cells, while lamin A/C was bound to these regions in LCL cells only (**Supplementary Figure 2**). Overall, these results indicate that viral LADs are present in the EBV genome and are enriched for lamin B1 in Type I latency and for lamin A/C in Type III latency.

**Figure 4:**
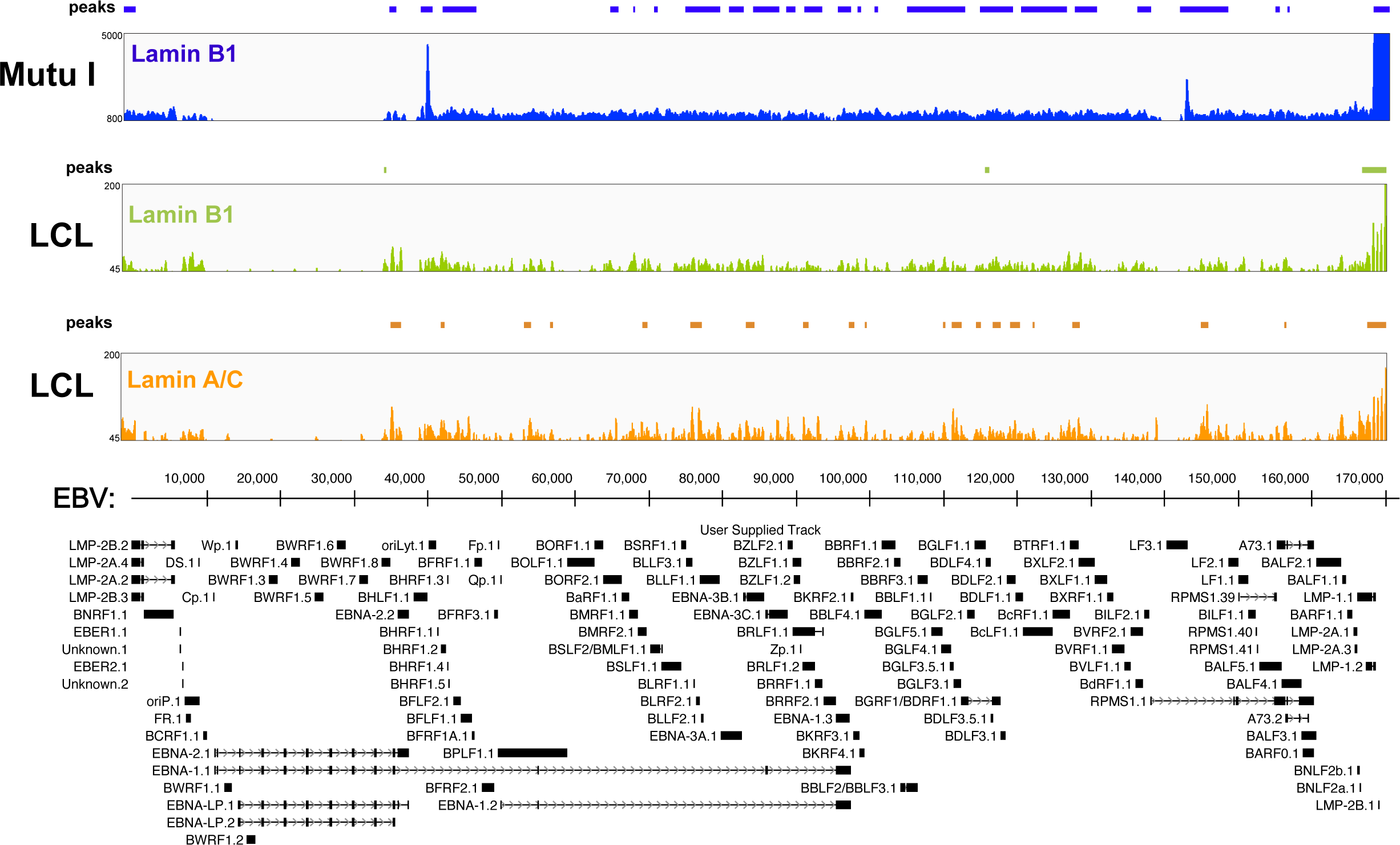
The EBV genome interacts with the nuclear lamins. **A)** ChIP-Seq analysis of lamin B1 and lamin A/C binding in EBV+ B cells adopting Type I (Mutu I) or Type III (LCLs) latency. Blue track profile: lamin B1 binding in type I cells; green track profile: lamin B1 binding in type III cells; orange track profile: lamin A/C binding in type III cells. The peaks identified through peak calling method are indicated above each track. Tracks are aligned with the annotated EBV genome shown at the bottom.

### Different pattern of lamina association between Type I and Type III EBV genomes

The differences in expression of lamin A/C between Type I and Type III latency and the presence of lamin A/C in the nucleoplasm of infected cells prompted us to study the relationship between lamin A/C and lamin B binding across the EBV genome in Type I and Type III latency. We applied a Spearman correlation using the number of reads at each peak as variables (**Supplementary Figure 3**). We found that lamin B1 binding strongly correlated between Mutu I and LCL cells and with lamin A/C binding in LCL cells (**Supplementary Figure 3**). Next, we generated an UpSet plot [59] to visually quantify the relationship between lamin B1 and lamin A/C binding across the EBV genome in Mutu I and LCLs. We observed similarity and differences between latency types with respect to the nuclear lamina interactions. We confirmed lamin B1 binding to three regions both in Mutu I and LCL cells. These regions also interacted with lamin A/C in LCL cells (**Figure 5A** and **5B**). In contrast, 8 LADs were bound by lamin B1 in Mutu I cells and lamin A/C in LCL cells, suggesting that the transition between latency types is associated with a switch in nuclear lamina components at these LADs (**Figure 5A**), which include the region downstream the lytic promoter Zp (∼ 90 Kk) and the region of RPMS-OriLyt (140 -150 Kb) (**Figure 5C**). Finally, we identified 12 regions that exclusively interacted with lamin B1 in Mutu I cells and 5 regions that exclusively interacted with lamin A/C in LCL cells (**Figures 5A** and **5D**). Overall, the viral LADs formed by lamin B1 and lamin A/C in the two different EBV latency types partly overlap, however, some LADs contain only one of the two nuclear lamina components. This indicates a rearrangement of the nuclear lamina-EBV genome interactions between latency types and suggests that the nuclear lamina may play alternative functions in regulating EBV gene expression.

**Figure 5:**
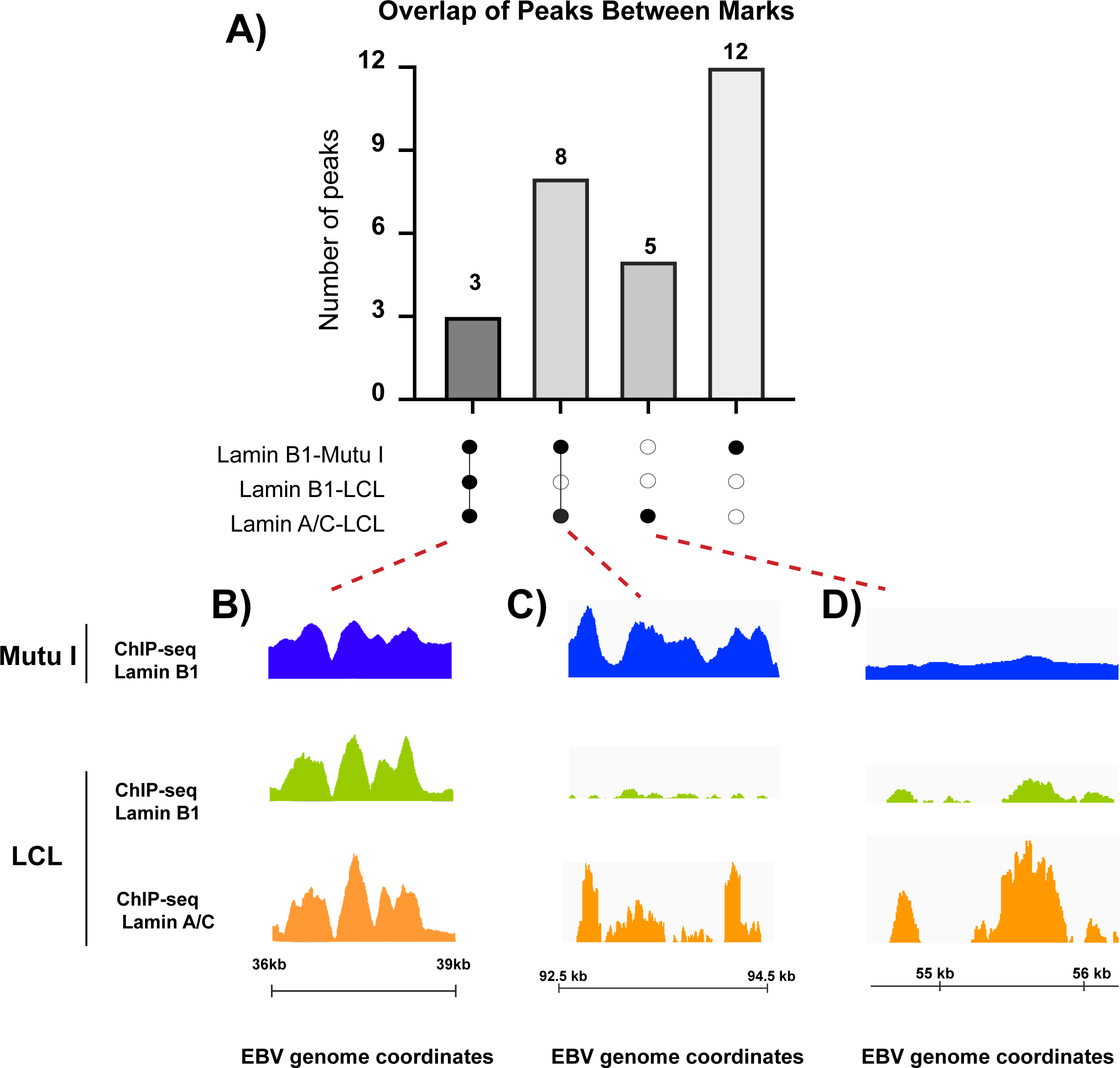
Patterns of lamin B1 and lamin A/C binding on the EBV genome in different types of latency. **A)** Upset plot of EBV peaks identified in three ChIP-seq datasets. Each bar in the upper plot corresponds to the number of peaks identified in the datasets indicated by black dots in the lower plot. Connected dots indicate an intersection of peaks between ChIP-Seq datasets. The number of EBV regions in each condition is indicated above the corresponding columns. **B), C) and D)** Lamin B1 and lamin A/C ChIP-Seq tracks for unique and overlapping peaks across the viral genome identified in A: LADs with lamin B1 binding in Mutu I and LCL cells and lamin A/C binding in LCL cells (B); LADs with lamin B1 binding in Mutu I cells and lamin A/C binding in LCL cells (C); LADs with lamin B1 binding only in Mutu I cells and lamin A/C binding only in LCL cells (D).

### Lamin A/C binding regulates the chromatin composition of the EBV genome

Since lamin B1 and lamin A/C binding regulates gene expression, our data suggest that the nuclear lamina proteins may regulate EBV latency. To test this hypothesis, we used CRISPR/Cas9 gene editing to knock out lamin A/C expression and assess the effect of its depletion in LCL cells (**Figure 6**). We focused on lamin A/C because its expression is linked to EBV infection and specific to type III latency. Furthermore, B-type lamins, including lamin B1, are essential for mammalian cells [60, 61], therefore their depletion in B cells may lead to cell death. We used two different single-guide RNAs (sgRNAs) that target distinct regions of the *LMNA* gene. After puromycin selection, we obtained two stable LCL cell lines in which we successfully knocked out *LMNA* gene expression, as verified by western blot and immunofluorescence analysis (**Figure 6A** and **6B**). We found that lamin A/C depletion had no effect on lamin B1 expression and localization and did not alter the nuclear shape (**Figure 6B**). We then tested whether the interaction of lamin B1 with the EBV genome was affected in lamin A/C KO cells. We used quantitative ChIP to quantify lamin A/C and lamin B1 binding at the OriLyt-left and TR regions, two of the viral LADs bound by both nuclear lamin proteins in type III LCL cells and where prominent lamin B1 peaks were observed in type I Mutu cells (**Figure 4** and **Figure 5B**). In lamin A/C KO cells, lamin B1 binding increased at OriLyt-left and decreased at the TR regions (**Figure 6C**), suggesting that the effect of lamin A/C on the binding of the nuclear lamina components may vary between different regions of the EBV genome.

**Figure 6:**
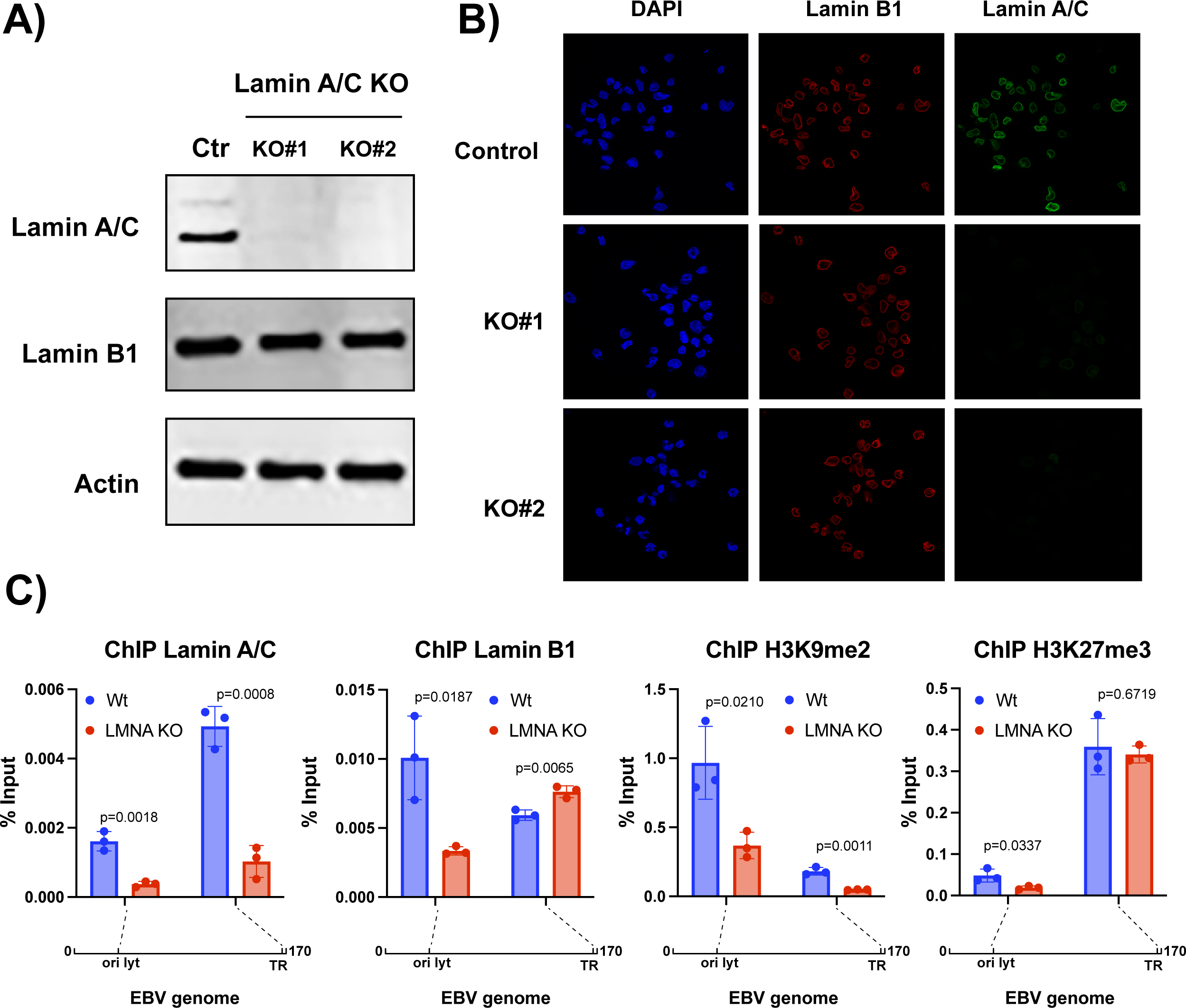
Lamin A/C knockout alters interaction of the EBV genome with the nuclear lamina and changes viral chromatin composition. **A)** Western blot analysis for lamin A/C, lamin B1 and actin protein expression in Ctr and *LMNA* KO GM12878 LCL cells. CRISPR/Cas9 gene editing was used to knock out lamin A/C expression in GM12878 cell lines using two different single-guide RNAs (sgRNAs) that target distinct regions of the *LMNA* gene. We generated two lamin A/C KO stable bulk populations. B) Immunofluorescence confocal microscopy analysis of Ctr and KO cells immunostained with lamin A/C (green) and lamin B1 (red) antibodies and DAPI (blue). C) Quantitative chromatin immunoprecipitation (ChIP-qPCR) analysis of EBV chromatin extracted from Ctr and *LMNA* KO GM12878 cells. EBV chromatin was analyzed for the binding of lamin B1 and lamin A/C and the deposition of the repressive histone marks H3K9me2 and H3K27me3 at the indicated regions. Data are presented as %input. N=3, Mean ± SD. The t test p values for the KO/ctr comparison are indicated.

Chromatin regions that interact with the nuclear lamina are enriched for specific histone marks: H3K9me2 [29]—a marker associated with localization at the nuclear periphery, and H3K27me3 [36]—a marker associated with heterochromatin and downregulation of gene expression. We assessed the effect of lamin A/C on H3K9me2 and H3K27me3 deposition at OriLyt-left and TR regions and found that in KO cells, H3K9me2 deposition was reduced at both viral sites, while H3K27me3 deposition was reduced only at OriLyt-left (**Figure 6C**), consistent with the observed differences in lamin B1 binding between these two regions in KO cells.

In mammalian cells, the LADs borders are demarcated by binding of CTCF [36], a protein implicated in chromatin loop formation. We recently reported that CTCF regulates changes in the three-dimensional structure of the EBV genome between types of latency [22]. This prompted us to investigate the relationship between lamin A/C and chromatin looping across the EBV genome. We used *in situ* HiC assay followed by enrichment for EBV-EBV chromatin interaction as we recently described [22], to assess the 3D structure of the EBV genome in lamin A/C KO cells and Ctrl cells (**Supplementary Figure 4**). We found few loops with a fold change ≥ 2 and with a p value < 0.05%. (**Supplementary figure 4A)**. When we corrected our analysis for multiple testing, we observed no loops with a false discovery rate (FDR) <5%, indicating that lamin A/C depletion did not cause significant changes in the EBV chromatin architecture (**Supplemental Figure 4B**). Taken together, these findings demonstrate that lamin A/C binding during latency influences the interaction of the EBV genome with the other component of the nuclear lamina and regulates viral chromatin composition without affecting the chromatin architecture.

### Lamin A/C regulates EBV gene expression during latency

We and others have demonstrated that epigenetic modifications play an essential role in regulating viral gene expression during latency. In lamin A/C KO cells, we observed changes in both lamin B1 occupancy and H3K9me2 and H3K27me3 deposition at viral LADs. To test whether EBV gene expression was altered as a consequence of lamin A/C depletion, we used RNA-Seq to analyze EBV global gene expression in KO and Ctr LCL cells (**Figure 7**). We performed an unsupervised principal component analysis (PCA) to assess the dominant direction of viral gene expression variability in our RNA-Seq datasets (**Figure 7A**). We observed that the first PC is associated with lamin A/C KO, with 61% of the viral gene expression variance between samples explained by the biological absence of lamin A/C (**Figure 7A**). We identified 27 EBV genes differentially expressed (p<0.05) between Ctr and lamin A/C KO cells, corresponding to a change in the expression of ∼25% of all EBV genes (**Figure 7B**). Through an unsupervised hierarchical cluster analysis, we found that all of the 11 EBV genes that were downregulated in lamin A/C KO cells encoded proteins that play a role during viral lytic replication (**Figure 7B**), while 14 of the 16 upregulated genes encoded EBV latent proteins, including EBNA3A, -3C, - 3D, LMP1 and LMP2, the EBER family, and EBNA2 (**Figure 7B**). The transcriptome profile of lamin A/C KO LCL cells is consistent with a role of lamin A/C in downregulating Type III latency and supporting lytic gene expression. Overall, our results show that the nuclear lamina is an important regulator of EBV viral gene expression during latency.

**Figure7:**
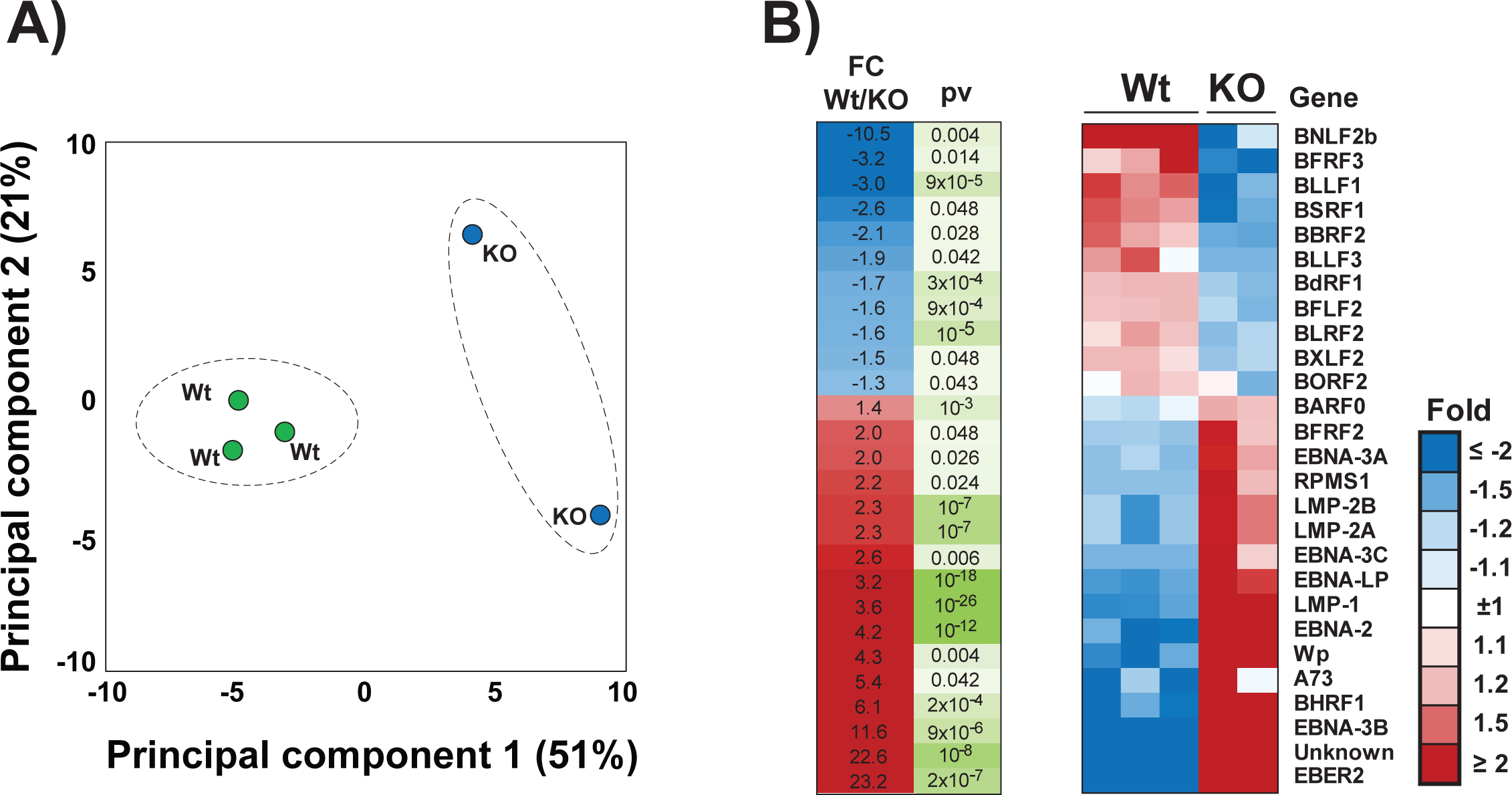
Lamin A/C knockout deregulates EBV gene expression in Type III B cells. **A)** Principal Component Analysis (PCA) of RNA-Seq analysis of Ctr and KO LCL cells (3 samples and 2 samples, respectively). The samples are shown as a function of principal component 1, or PC1 (x-axis), and principal component 2, or PC2 (y-axis). The percentage of variance explained by PC1 and PC2 is indicated. **B)** Heat map of RNA-seq data from Ctr and *LMNA* KO LCL cells showing genes whose expression was significantly altered (P<0.05) by lamin A/C depletion, which include EBNAs and LMP1 family members; differences were calculated using p-values.

## DISCUSSION

EBV latent infection is characterized by alternative and dynamic expression patterns of viral latent genes, which correspond to distinct latency types. These latency types exert different effects on infected cells and correlate with different EBV malignancies. We and others have demonstrated that the EBV latency types are associated with different levels of DNA methylation and histone modifications across the viral genome, indicating the importance of epigenetics in establishing and maintaining EBV latency [18, 62]. How the viral epigenetic patterns are regulated is only partially understood. The nuclear lamina is an important regulator of the epigenome as it interacts with specific regions of the chromatin and influences its composition. A role of the nuclear lamina in regulating herpesviruses infection, including EBV, has been reported during viral lytic activation, but, to the best of our knowledge, the ability of the nuclear lamina to regulate EBV gene expression during latency has been unexplored. Here, we report that EBV latent infection alters the composition of the nuclear lamina and is linked to lamin A/C expression. We found that Type III latent EBV+ B cells express lamin A/C in addition to lamin B1 and observed similar changes in lamin A/C expression upon EBV primary infection, which triggers germinal center-like differentiation of naïve B cells, consistent with lamin A/C expression being correlated with cell differentiation [63, 64]. Our temporal proteomic and transcriptomic analyses reveal indeed a steady increase in lamin A/C expression starting 4 days after EBV infection, when the first cell division is observed in infected B cells [58].

EBV infection of primary B cells triggers cell activation and proliferation by mimicking fundamental B cell signaling pathways, including the BCR and CD40 pathways. Our results show that CD40 signaling triggers lamin A/C expression, confirming previous reports that in immune cells, lamin A/C is rapidly expressed upon antigen activation [39]. Since the EBV protein LMP1 mimics CD40-mediated B cell activation [65, 66], we speculated that lamin A/C expression may be dependent on LMP1. Interestingly, work from the Gewurz laboratory showed that LMP1 is expressed 4 days post infection of primary B cells, which temporally coincides with expression of lamin A/C in EBV-infected B cells in our analysis [57]. Overall, our results indicate that the induction of lamin A/C is part of the transcriptional reprogramming induced by EBV infection to mimic CD40 stimulation and trigger germ-like differentiation of naïve B cells.

The changes in nuclear lamina composition observed upon EBV infection of B cells and between Type I and Type III EBV+ cell lines, and the well-established link between nuclear lamina and gene regulation, prompted us to investigate whether lamin B1 and lamin A/C interact with the EBV genome during latency. We report for the first time the existence of LADs across the EBV genome. We found that the pattern of lamin B1 and lamin A/C binding to the EBV LADs varies between EBV latency types. Our data show that in Type I latency, lamin B1 extensively binds the EBV genome at 23 regions. Since chromatin regions that interact with the nuclear lamina are repressed, this observation is concordant with the robust transcriptional repression of viral genes in Type I latency and suggests that transcriptionally silent EBV regions may localize at the nuclear periphery, where lamin B1 is located. Notably, in EBV+ gastric cancer cells, in which viral gene expression is limited, genome-wide associations between the EBV chromosome and the host genome tend to occur at genomic regions associated with the nuclear lamina [67]. During Type III latency, lamin B1 binding was restricted to 3 regions, while lamin A/C binding was prevalent (16 regions). Among the regions where lamin B1 binding was conserved in both latency types and overlapped with lamin A/C binding, we observed the two origins of lytic replication OriLyt-left and RMPSI-OriLyt that are inactive during latency, pointing to a role of the nuclear lamina in regulating viral lytic genome replication. These observations are consistent with work from the Miranda group showing that upon EBV lytic reactivation, when both OriLyt regions are active, the interactions between EBV and human LADs decrease [68]. Although lamin B1 and lamin A/C colocalized at the nuclear rim, a lamin A/C nucleoplasmic pool was present in type III EBV+ cells, suggesting it could interact with the chromatin. We speculate that in Type III latency the EBV regions exclusively associated with lamin A/C may be localized in the proximity of the nuclear lamina but in a less peripheral position than lamin B1-associated viral regions in Type I latency. This hypothesis is supported by the observation that in LCL and Raji cells, which adopt a Type III latency, the EBV genome is localized at the perichromatic region of the nucleus [69]. Overall, our results uncover the dynamic interaction existing between the EBV genome and lamins during latency and reveal the importance of lamin A/C expression in repositioning the EBV genome with respect to the nuclear periphery.

Rearrangement of the lamin-genome interactions and changes in lamin composition of the LADs contribute to changes in gene expression. We observed that lamin A/C depletion alters the expression of 27 of the ∼ 100 genes expressed by EBV and exerts specific and opposite effects on the expression of latent and lytic genes. In fact, depletion of lamin A/C resulted in upregulation of latent genes and downregulation of lytic genes, suggesting a role of lamin A/C in fine-tuning EBV gene expression during latency. Our data suggest therefore that EBV infection induces expression of lamin A/C, which subsequently binds the viral genome and modulates the expression of viral latent genes. This hypothesis is consistent with published transcriptional profiling of EBV infection of B cells showing that in the pre-latent viral phase of the infection, the peak of lamin A/C expression coincides with reduced expression of EBNA2, BHRF1, and EBNA-LP [58]. More work will be necessary to better understand the role of lamin A/C during the early stages of EBV infection, but our data establish a role of this protein in transcriptionally regulating viral gene expression during latency.

The nuclear lamina-genome interactions regulate gene expression by modifying the chromatin composition (i.e., histone modifications). In Type III latency, we observed that lamin A/C depletion resulted in reduced H3K9me2 deposition at viral loci bound by both lamin A/C and lamin B1. The H3K9me2 histone mark is specifically associated with heterochromatin regions positioned at the nuclear periphery, and reduced H3K9me2 levels at LADs weaken the association of these regions with the nuclear lamina [29]. Our results therefore highlight the role of the nuclear lamina in regulating the chromatin composition of the EBV genome by affecting H3K9me2 deposition across the viral genome during latency. We hypothesize that this altered histone modification may play an important and underappreciated role in regulating EBV latency. We plan to further explore the role of the H3K9me2 histone mark in regulating the EBV epigenome and viral gene expression by inhibiting the H3K9 methyltransferase G9a.

Lamin A/C and lamin B1 binding regulates the three-dimensional organization of the cellular genome, and our recent work has showed the importance of 3D chromatin structure in controlling viral gene expression between latency types [22]. However, we did not observe effects on the 3D-strucuture of the EBV episome after lamin A/C depletion, indicating that lamin A/C binding changes the local composition of the viral chromatin without affecting its architecture.

Based on our observations on the location of lamin A/C within the nuclear space, we propose that upon EBV infection, lamin A/C accumulates in the nucleus of infected cells, where it binds the EBV genome and repositions it toward the nuclear periphery, attenuating viral gene expression. As lamin A/C is present both at the nuclear periphery and in the nuclear interior, the association of lamin A/C EBV LADs with the nuclear lamina is dynamical, allowing expression of a few viral latent genes. Rearrangement of the nuclear lamina-EBV genome interactions and the observed lamin B1/lamin A/C switch may result in heterochromatin deposition across the viral genome and repression of EBV gene expression. How remodeling of the viral LADs occurs remains unknow, but since B cell differentiation is known to induce a reorganization of the nuclear space, we argue that the effects of EBV infection on B cell differentiation may induce this remodeling.

Further studies are required to validate our hypothesis, but our results highlight for the first time the dynamic interplay between lamin A/C and lamin B1 and the EBV genome during viral latency and the regulatory role of the lamin-genome interaction in restricting viral latent gene expression.

The nuclear lamina plays an anti-viral role in herpesvirus replication, including EBV, HSV1 and HSV2, and HCMV, representing a physical barrier that prevents viral capsid egress. Elegant work from the Knipe group also showed the importance of the nuclear lamina in the formation of HSV-1 viral replication compartments [43–49]. Our data shed light on an additional antiviral function during EBV latency, by contributing to the formation of an epigenetic “barrier” across the viral genome that restricts viral gene transcription.

Our work has some limitations. It is possible that lamin A/C may work as a repressor of latent gene expression in cooperation with other cellular factors or transcription factors that can help tether the EBV viral genome at the nuclear periphery. It is worth noting that a similar model has been proposed for the regulation of HSV-1 lytic gene expression by lamin A/C [46]. It will be of interest to identify the cellular factor(s) that cooperate with the nuclear lamina to anchor the EBV genome at the nuclear periphery. The changes we observed in the composition of the nuclear lamina may also affect host gene expression, an aspect that we left unexplored. Although such studies are beyond the scope of this work, the role of lamin A/C regulating host gene expression may be relevant to the creation of a cellular environment that supports Type III viral latency and EBV-induced cell proliferation.

In summary, our results, together with previous work from our group and others, reveal a novel and unrecognized function of the nuclear lamina in the host response against the intrusion of viral DNA into the nucleus. Our work adds an important piece of information to the existing model of how host epigenetic factors regulate the viral chromatin, and how host and viruses communicate to regulate gene expression through chromatin modification. Gaining a better insight into the nuclear lamina/EBV interaction will help reveal additional molecular steps that are necessary for cells to control viral replication.

## MATERIALS AND METHODS

### Cell culture and treatment

Cell lines were maintained in a humidified atmosphere containing 5% CO_2_ at 37°C. EBV positive cell lines were cultured in suspension in RPMI 1640 supplemented with fetal bovine serum at a concentration of either 10% for type I latency (Mutu and Kem I) or 15% for type III latency (LCL, Kem III, and GM12878). All cell media were supplemented with 1% penicillin-streptomycin.

### CRISPR Cas9 Mutagenesis

B-cells were transduced with lentiviruses expressing sgRNAs, as previously described [70]. Briefly, lentiviruses were produced by transfection of 293T cells, which were plated in a 6-well dish at a density of 600,000 cells per well in 2 mL DMEM supplemented with 10% fetal calf serum 24 hours before transfection. Transfection was done using the TransIT-LT1 Transfection Reagent (Mirus). Two solutions were prepared for each well: a solution with 4 mL of LT1 diluted in 16 mL of Opti-MEM (Corning) and incubated at room temperature for 5 min. The second solution contained 150 ng pCMV-VSVG (Addgene 8454), 400 ng psPAX2 (Addgene 12260), and 500 ng plentiguide puro expression vector (Addgene cat # 52963), in a final volume of 20 mL with Opti-MEM. The two solutions were then mixed and incubated at room temperature for 30 min, added dropwise to wells and gently mixed. Plates were returned to a 37C humidified chamber with 5% CO2. The following day, media was exchanged to RPMI with 10% fetal calf serum. Virus supernatant was harvested 48h post-transfection and filtered through a 0.4 uM sterile filter. Fresh media was replenished on the 293T producer cell and a, second harvest, was done 72h post-transfection and again sterile filtered. Virus supernatant was added to target cells at 48- and 72-hours post-transfection. Target cell selection was begun 48 hours post-transduction by addition of puromycin 3g/ml. sgRNA sequences used were as follows. Control: ATTTCGCAGATCATCGACAT. LMNA: GCGCCGTCATGAGACCCGAC and AGTTTAAGGAGCTGAAAGCG.

### B cell activation and B cells infection

Platelet-depleted venous blood, obtained from the Brigham and Women’s Hospital bank, were used for primary human B cell isolation, following institutional guidelines for discarded and de-identified samples. CD19+ B-cells were isolated by negative selection as follows. RosetteSep (Stem Cell Technologies, #15064) and EasySep negative isolation kits (Stem Cell Technologies, #19054) were used to sequentially isolate B-cells, with the following modifications of the manufacturer’s protocols. For RosetteSep, 40 μL of antibody cocktail was added per mL of blood and layered onto Lymphoprep density medium for centrifugation. For EasySep, 10 μL of antibody cocktail was added per mL of B cells, followed by 15 μL of magnetic bead suspension per mL of B cells. Following this negative selection, FACS for CD19 plasma membrane expression was used to confirm B cell purity. The following reagents were used for B-cell stimulation: MEGA-CD40L (Enzo Life Sciences Cat# ALX-522-110-C010, 50ng/mL), CpG (IDT, TCGTCGTTTTGTCGTTTTGTCGTT, 1μM), αIgM (Sigma Cat#10759, 1mg/mL), IL-4 (R&D Systems Cat#204-IL-050, 20ng/mL), αIgG (Agilent Cat#A042402-2). Cells were cultured in RPMI-1640 (Invitrogen), supplemented with 10% standard FBS and penicillin-streptomycin. Cells were cultured in a humidified incubator at 37 C and at 5% CO2.

### Western blot analysis

Cell lysates were prepared in radioimmunoprecipitation assay (RIPA) lysis buffer (50 mM Tris-HCl, pH 7.4, 150 mM NaCl, 0.25% deoxycholic acid, 1% NP-40, 1 mM EDTA; Millipore) supplemented with 1× protease inhibitor cocktail (Thermo Scientific). Protein extracts were obtained by centrifugation at 3,000 × *g* for 10 min at 4°C. Protein concentration was measured using a bicinchoninic acid (BCA) protein assay (Pierce). Lysates were boiled with 2× Laemmli sample buffer (Bio-Rad) containing 2.5% β-mercaptoethanol (Sigma-Aldrich). Proteins were resolved by gel electrophoresis on a 4 to 20% polyacrylamide gradient Mini-Protean TGX precast gel (Bio-Rad) and transferred to an Immobilon-P membrane (Millipore). Membranes were blocked in 5% milk in PBS-T for 1 h at room temperature and incubated overnight at 4°C with primary antibodies against Lamin AC (Active Motif 39287), Lamin B1 (Abcam ab16048), EBNA2 (Abcam ab90543), LMP1 (Abcam ab78113), and actin (Sigma-Aldrich A2066) as recommended per the manufacturer. Membranes were washed, incubated for 1 h with the appropriate secondary antibody, either goat anti-rabbit IgG-HRP (Jackson Immuno Research) or rabbit anti-mouse IgG-HRP (Jackson Immuno Research). Membranes were then washed and detected by enhanced chemiluminescence.

### Proteomic analysis

Samples were prepared for LC-MS/MS experiments are described previously [71]. Samples were prepared for LC-MS/MS analysis as described above except samples were normalized to total protein amount. A protein bicinchoninic acid assay was used on clarified lysate before reduction and alkylation. Additionally, after labelling samples with TMT-11 reagents, samples were normalized based on ratio check data to ensure a 1:1:1:1:1:1:1:1:1 total protein ratio was observed as described previously [71]. For each BPRP fractionated sample, 12 of the 24 fractions were selected for LC-MS/MS analysis to reduce redundant identifications. Each fraction was analyzed on a Thermo Orbitrap Fusion Lumos with a Proxeon EASY-nLC 1200 system before the source (Thermo Fisher Scientific, San Jose). The mass spectrometer was operated in a data-dependent centroid mode for all SPS-MS3 methods. On-line chromatography was performed on a 100 μm inner diameter microcapillary column packed with 35cm of Accucore C18 resin was used. Approximately 2μg of labelled peptides were loaded onto the column. Spectra were acquired across a 90-minute LC gradient ranging from 6-25% acetonitrile in 0.125% formic acid. For the 10-condition experiment, a real-time search method was employed in the instrument such that SPS-MS3 scans would not trigger unless a successful Comet search result was obtained from a preceding MS2 scan [72]. Spectra from all mass spectrometry experiments were processed and searched with an in-house Sequest-based software pipeline as described previously [71, 73, 74]. Raw data acquired on mass spectrometer instruments were converted into an mzXML format before a Sequest search was executed against a human proteome database (Uniprot Database ID: 9606, downloaded February 4, 2014). This database was concatenated with common laboratory contaminants and a database comprised of all protein sequences reversed. Ion tolerances for precursor and product ions were set to 50ppm and 0.9 Da respectively. A variable mass difference of +15.99491 Da was assigned to methionine residues to account for potential oxidation. Additionally, fixed modifications on cysteine (+57.02146 Da) and lysine and the peptide N-terminus (+229.16293 Da) were assigned to account for protective alkylation and TMT-11 labelling, respectively.

### RNA extraction and RNA-seq

Total RNA for Lamin A/C knockout experiment was isolated from 1.5 x 10^6^ cells using using a PureLink RNA Mini Kit (ThermoFisher) according to the manufacturer’s protocol. RNA samples were submitted to the Wistar Institute genomics core facility for initial analysis of RNA quality, with each sample having a RIN value greater than 8.5 (TapeStation, Agilent Technologies). Sequencing library preparation was then completed using the QuantSeq 3’-mRNA kit (Lexogen) to generate Illumina-compatible sequencing libraries according to the manufacturer’s instructions. Single reads of 75 bp were obtained using a NextSeq 500 sequencer. RNA-seq data was aligned using bowtie2 [75] against hg19 version of the human genome and all unaligned reads were then aligned against NC_007605.1 version of EBV genome and RSEM v1.2.12 software [76] was used to estimate raw read counts and RPKM for EBV genes. DESeq2 [77] was used to estimate significance of differential expression between groups pairs. Genes that passed nominal p<0.05 (FDR<5%) threshold were reported.

### Immunofluorescence assay

For cell seeding sterile cover slips or slides can be used and coated with adhesive material, Poly-L-Lysine (PLL). ∼500 ul of 0.5 *10^6/ml cells resuspended in media can be seeded on the coverslip and incubate overnight in a humidified atmosphere containing 5% CO_2_ at 37°C. Cells were fixed with 4% PFA and incubate at room temperature for 10 minutes. Cover slips washed for three times with Ca2+/Mg2+ free PBS. Then permeabilize in PBS + 0.1% Triton X for 10 minutes at room temperature and then washed for three times with Ca2+/Mg2+ free PBS. Coverslip were blocked in chi block (5% serum, 1%BSA in Ca2+/Mg2+ free PBS) for 1 hour at room temperature. Incubation with primary antibody diluted in chi block [78]at 1:100 at room temperature for an hour. Wash with PBS + 0.1% Triton X for three times, each wash 5 minutes. Incubation with secondary antibody, diluted 1:1000 in chi block and incubated at room temperature for an hour. Wash with PBS + 0.1% Triton X for three times, each wash 5 minutes last wash without tween. Mounting by adding a drop of pro Long Diamond antifade mountant with DAPI, let it mount overnight and image the next day.

### ChIP-seq

Chromatin immunoprecipitation with next-generation sequencing (ChIP-seq) was performed as previously described [78] with minor changes. A total of 2.5 × 10^7^ cells were cross-linked with 1% formaldehyde for 10 min, and chromatin was sonicated with a Qsonica sonicator. One-tenth of the sonicated chromatin was collected and used as input material, while the rest of the chromatin was immunoprecipitated using 5 μg of Lamin AC antibody (Active Motif), 5 μg of Lamin B1 antibody (Abcam), or in LCLs and Mutu I. DNA fragments of ~150 to 300 bp were visualized by agarose gel purification. Immunoprecipitated DNA was ligated to adapter primers using the TruSeq ChIP library preparation kit (Illumina) and then sequenced using the Illumina HiSeq 2500 platform according to the manufacturer’s recommendations (Illumina) at the Fox Chase Cancer Center Sequencing Facility. For both LCL and Mutu I, ∼15 ng of Lamin A/C, and B1-immunoprecipitated or input DNA was recovered from each biological replicate. Sequenced reads were trimmed for quality and sequencing adapters using trimmomatic [79] then aligned to the EBV genome using bowtie2 [75]. Aligned reads were prepared for visualization in IGV browser using samtools [80] and BigWig tools [81]. Peaks were called using macs2 broad peaks and differential peaks identified using macs2 bdgdiff [82]. Heatmaps and traces were made using deepTools [83]

### ChIP-qPCR

ChIP assays were performed according to the Upstate Biotechnology, Inc., protocol as described previously, with minor modifications [78]. Briefly, cells were fixed in 1% formaldehyde for 15 min, and DNA was sonicated using a Qsonica sonicator to generate 200- to 500-bp fragments. Chromatin was immunoprecipitated with antibodies to LaminB1 (abcam), Lamin A/C (Active Motif) and H3K9me2 (abcam). Real-time PCR was performed with a master mix containing 1× Maxima SYBR green, 0.25 μM primers, and 1/50 of the ChIP DNA per well. Quantitative PCRs were carried out in triplicate using the ABI StepOnePlus PCR system. Data were analyzed by the ΔΔ*C_T_* method (where *C_T_* is threshold cycle) relative to DNA input and normalized to the IgG control.

### HiC assay

Hi-C assay was performed as previously described (Reference to Nat. Comm. Paper). Briefly, 5x106 cells per condition were collected for in situ Hi-C. Libraries of total ligation products were produced using Ultralow Library Systems V2 (Tecan Genomics, part no. 0344NB-32) as per manufacturer’s protocol. Purified libraries were then enriched for only EBV genome ligation products using myBaits enrichment kit as per manufacturer’s protocol. Both enriched and human libraries were sequenced using the Illumina MiSeq500 sequencing platform with paired-end 75bp read length. Briefly, 75-bp paired reads were separately aligned to the EBV genome (V01555.2) using Bowtie2 (version 2.2.9) with iterative alignment strategy71. Redundant paired reads derived from a PCR bias, reads aligned to repetitive sequences, and reads with low mapping quality (MapQ < 30) were removed. Reads potentially derived from self-ligation and undigested products were also discarded. EBV genome were divided into 5 kb windows with 1 kb sliding. Raw contact matrices were constructed by counting paired reads assigned to two 5 kb windows. Hi-C biases in contact matrices were corrected using the ICE method. The ICE normalization was repeated 30 times. Significant associations were determined based on the distance between two 5kb windows, all combinations were categorized into 20 groups. We assumed Hi-C score as Poisson distribution with a parameter λ matching the mean score. We then assigned a P values for each group and applied an FDR correction for multiply hypotheses. FDR < 0.05 were defined as significant 27 associations. Significant associations were plotted as circos graph using the circlize package (version 0.3.3) of R (version 3.6.1). The detailed protocol with all minor alterations will be happily supplied by corresponding author per request.

### qRT-PCR

For quantitative reverse transcription-PCR (qRT-PCR), RNA was extracted from 2 × 10^6^ cells using TRIzol (Thermo Fisher Scientific) according to the manufacturer’s instructions. SuperScript IV reverse transcriptase (Invitrogen) was used to generate randomly primed cDNA from 1 μg of total RNA. A 50-ng cDNA sample was analyzed in triplicate by quantitative PCR using the ABI StepOnePlus system. Data were analyzed by the ΔΔ*C_T_* method relative to expression of 18S and normalized to controls.

### Histone Extraction

Cell were harvest and wash twice using ice-cold PBS. Cell were resuspended it in Triton Extraction Buffer (TEB: PBS containing 0.5% Triton X 100 (v/v), 2mM phenylmethyl sulfonyl fluoride (PMSF), 0.02% (w/v) NaN3 at a cell density of 107 cells per ml. Cell were lysed on ice for 10 minutes with gentle stirring, centrifugated at 2000 rpm for 10 minutes at 4 C. The supernatant was removed and discarded. Cells washed in half the volume of TEB and centrifuge at before. Pellets resuspend in 0.2N HCl at a cell density of 4x107 cells per ml. Acid extraction of histones over night at 4 C. Next day samples were centrifuge at 2000 rpm for 10 minutes at 4 C. The supernatant moved to a new clean tube and determinate the protein content by Bradford assay.

### Statistical analysis

All experiments presented were conducted at least in triplicate to ensure reproducibility of results. The Prism statistical software package (GraphPad) was used to identify statistically significant differences between experimental conditions and control samples, using Student’s *t* test as indicated in the figure legends.

## AKNWOLEDGMENTS

Research reported in this publication was supported by the National Institute of Allergy and Infectious Diseases of the National Institutes of Health under Award Number R01AI130209 to IT and R01AI164709 and CA228700 to BG, a Burroughs Wellcome Career Award in Medical Sciences, and a Lymphoma Research Foundation Fellowship to RG. We thank Paul Lieberman for scientific discussions the Wistar Imaging Facility for technical assistance.

---

**Supplementary Figure 1:** The graph shows the percentage of CD69+ and CD69-cells after FACS sorting of primary B cells treated for 24 hours with a stimulatory cocktail containing 20 ng/mL Interleukin 4 (IL-4), 5uM CD40 ligand (CD40L), and 25 ng/mL CpG oligodeoxynucleotides.

**Supplementary Figure 2:** Quantitative chromatin immunoprecipitation (ChIP-qPCR) analysis of EBV chromatin extracted from Mutu I and LCL EBV+ cells. EBV chromatin was analyzed for the binding of lamin B1 and lamin A/C at the indicated EBV regions. Data are presented as %input. N=3, Mean ± SD.

**Supplementary Figure 3:** Heat map of Spearman correlation coefficients of the read counts from all locations where a peak is present in any sample. The dendrogram indicates similarity between ChIp-seq samples based on read counts.

**Supplementary Figure 4: A)** Scatter plots of normalized HiC counts of DNA-DNA interactions across the EBV genome in Ctr (x-axis) and *LMNA* KO (y-axis) LCL cells. DNA-DNA interactions with p<0.05 are indicated in blue (downregulated in Ctr cells) and red (upregulated in KO cells). **B)** Circos graph of all DNA-DNA contacts across the EBV genome that change between Wt and *LMNA* KO LCL (GM12878) cells. DNA-DNA contacts derived from HiC matrices (chromatin loops) with a p<0.05 are shown. Blue arcs represent chromatin loops that are frequent in Ctr cells; red arcs represent chromatin loops that are more frequently observed in *LMNA* KO cells.

