## Supplementary figures and images for "The Nuclear Lamina Binds the EBV Genome During Latency and Regulates Viral Gene Expression"

### Supplemental figure 1

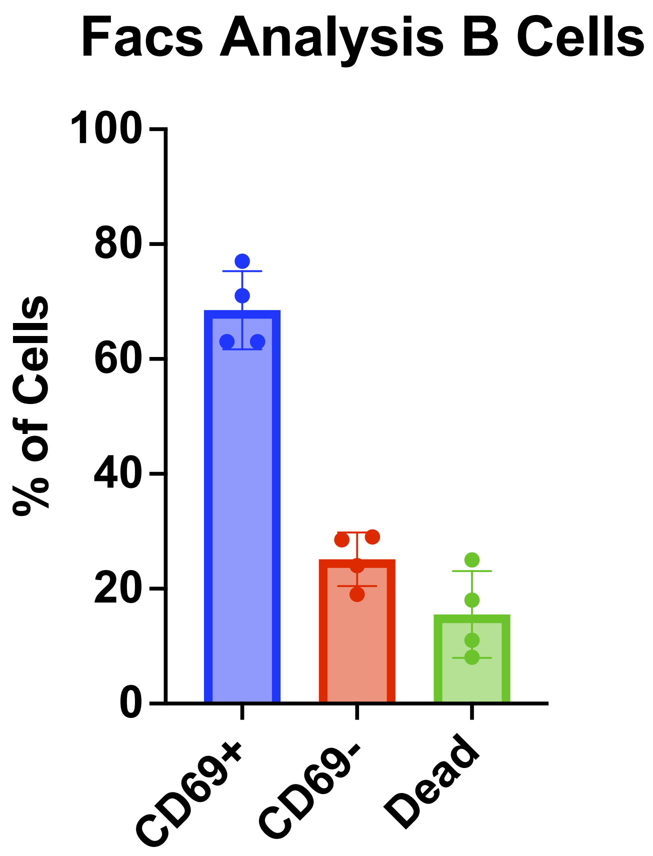

### Supplemental figure 2

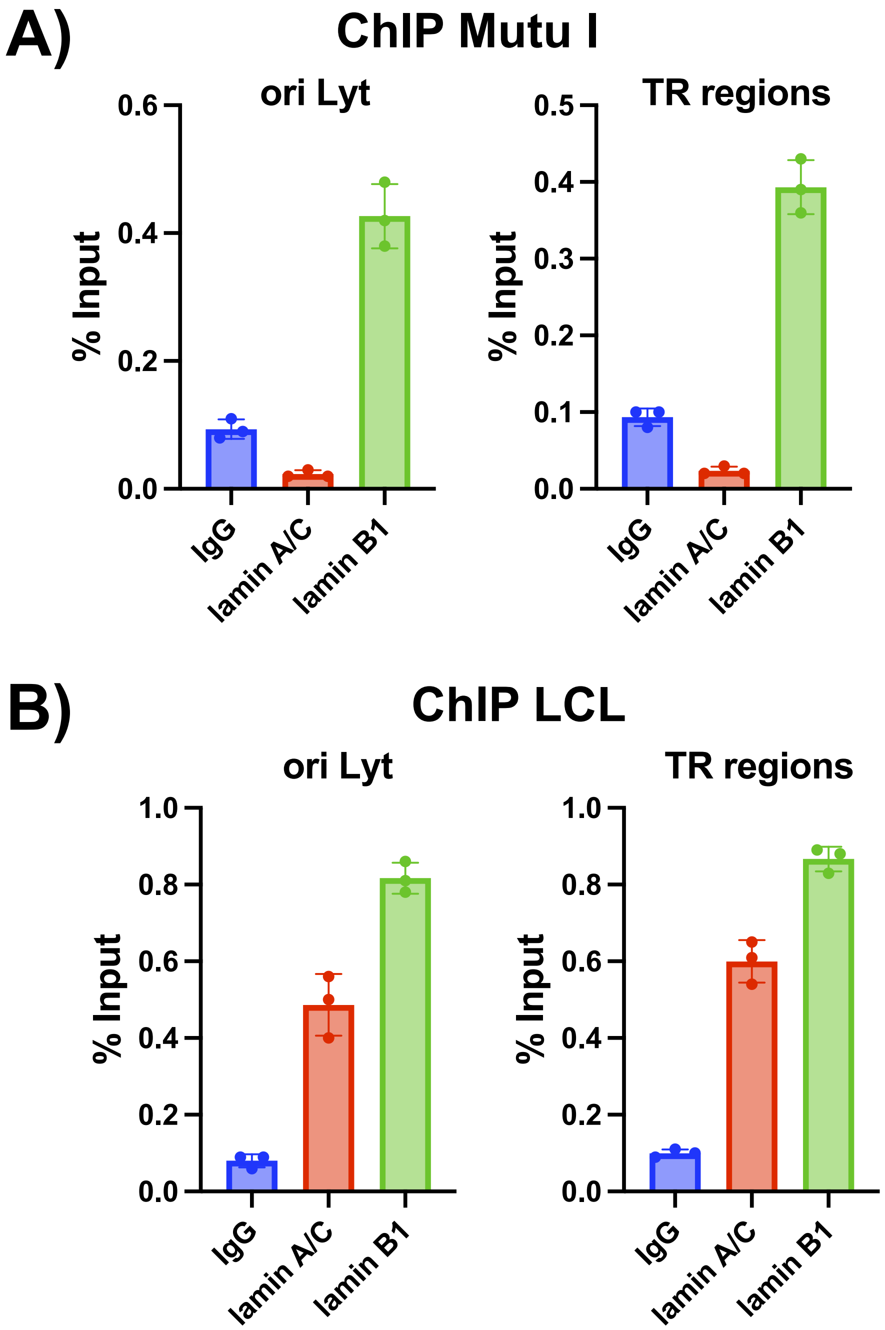

### Supplemental figure 3

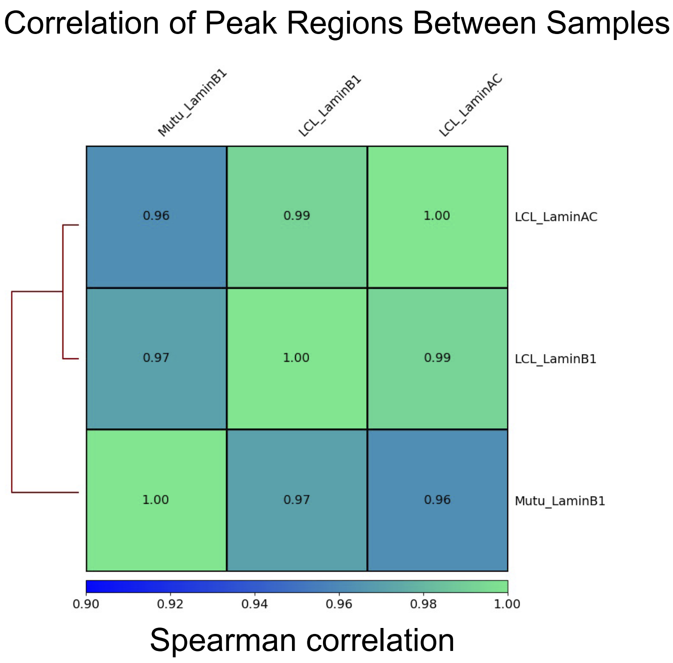

### Supplemental figure 4

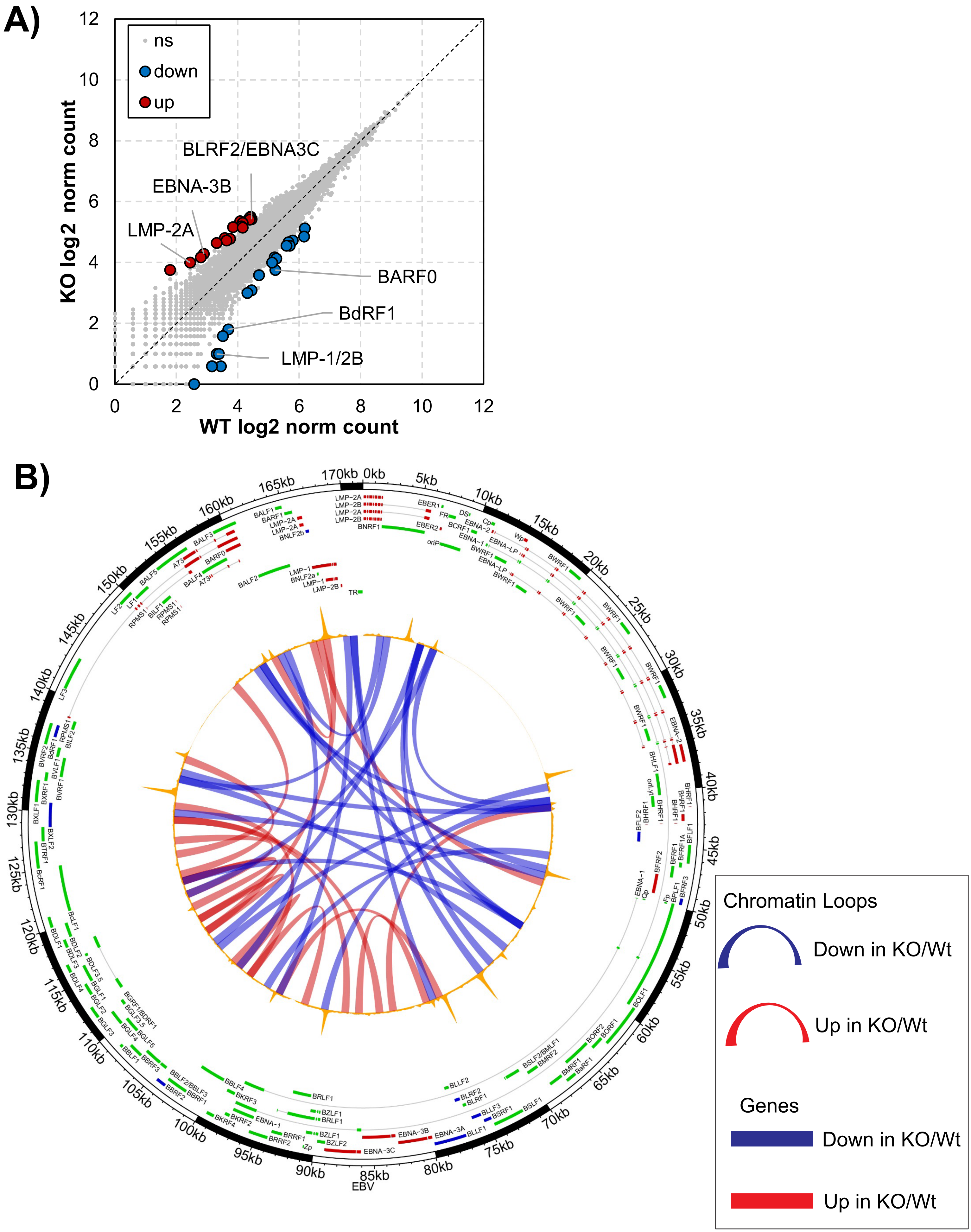
